# Properties and effect of Ittar in mice behavior and neurological functions

**DOI:** 10.1101/2019.12.15.876581

**Authors:** AbdulRahman Al Asmari, Faisal Kunnathodi, Riyasdeen Anvarbatcha, Mohammed Tanwir Athar, Abdul Quaiyoom Khan, Dhayasankar Sigmani, Mohammed M Idris

## Abstract

Fragrances are generally synthetic, natural or a combination of both. Ittar or Attar is a natural fragrance material synthesized from plants which were majorly used by people worldwide as a matter of pride, deodorization and attractant. The effect and role of using Ittar on mammalian system is still not known at the behavioral and neurological level. This study was aimed to understand the constituents and the effect of Ittar on mice by measuring their behavioral and neurological changes upon exposure to Ittar. Batches of animals were exposed to ten different types of Ittar for 1, 2 and 3 weeks respectively. Behavior of the animals exposed to the Ittar perfumes were analyzed against the control animals behavior for physiological activities. Also, the expression level of neurotransmitter such as Dopamine, 5-HIAA, HVA and Serotonin were evaluated based on HPLC analysis. Based on this study it was found that all the ten Ittars were different for its different constituents with some basic overlapping molecules among them. The neurotransmitter such as DOPA, 5-HT, Homovanillic Acid and Serotonin were found non-significantly increased in the total brain after a three week long continuous exposures of Ittar to the animal. Histological analysis of the ittar exposed mice skin showed mild to massive changes in both epidermal and dermal layers characterized with intense infiltration of inflammatory cells, hyperplasia, hyperchromasia and edema formation against control animals in few of the Ittars. From this study it is evident that Ittar increases mood of the animal through the up-regulation of various neurotransmitter but subtle effect on the skin because of the presence of various chemical constituents of the Ittar. Based on this study it is evident that use of Ittar as fragrance might be beneficial at the molecular level and it will be appropriate to apply without direct exposure to skin.

## Introduction

It has been long known that fragrances alter mood and brain chemistry. Fragrance plays an important role in inducing the pulse, mood and blood pressure in brain directly. It is well known that perfume induces a person’s love and mood with his opposite partner. A change in body odor leads to a phenomenal change in behavior of opposite partner. Comfort and discomfort of the partner is depending on the body odor and pheromones. Inhalation of fragrance leads to odor molecules to travel up the nose and get captured by specific receptor present in the olfactory epithelium. Olfactory bulb plays a big role in inducing the action of smell through the help of new neuron produced and migrated to olfactory bulb by neural stem cells. Granule cells, newborn neurons are inhibitory interneurons, play an important role in adaptation of new smells and odors [1]. Odor molecules stimulate the lining of nerve cells which trigger electrical impulses to the olfactory bulb, which transmits the impulses to the gustatory center, the amygdala and other part of the limbic system of the brain. Perfumes through their profound physiological and psychological effects directly controls the limbic system which controls heart rate, blood pressure, breathing, stress level, hormone balance and memory [2].

Perfumes are used as scent material towards masking the bad odor of our body secretion. Use of perfumes products causes exposure of scent material to skin, upper respiratory tract to lungs and olfactory pathway to the brain. Fragrance inhalation by the nose goes directly to the brain where its neurological effects can alter blood pressure, pulse and mood along with sedative effects [3]. As perfumes used are generally volatile and made from petroleum products, physical and neurological problems including headaches, dizziness, fatigue and dementia are associated with its use [3]. Volatile perfumes contain neurotoxins which have a causal link to central nervous system disorders, headaches, anxiety, depression and mood swings [4]. To avoid second hand exposures, hospitals and public place demands for not using perfumes.

Prior to the modern synthetic fragrance materials, people were using perfumes derived from flowers and plants. Floral based remedies have been used as a special way of treating patients [5]. It is also well known that aromas from natural sources influences body and mind. Fragrance from essential oils have claimed to enhance emotional state to lifespan. Natural fragrances and oils were been used for centuries as incense for religious and ritualistic purposes as it can influence the emotions and states of mind [6].

Ittar or Attar is a botanically derived natural perfume oil being used worldwide. The all-natural perfumes are prepared by either hydro or stream distilled from various part of the plants, mostly the flower part and natural sources. The alcohol-free perfume is being used both as a deodorant and an attractant for a long time. Ittar are classified in to several types based on types of distillation and source of extraction. No study has differentiated the effect of Ittar with the other perfumes. Also, there is no data available to understand why and what are the advantages and disadvantages of using the Ittar against the synthetic or chemically derived perfumes. In this study we aimed to analyze the effect of Ittar on mice behavior and neurological changes and its various constituents.

## Experimental

### Sample collection

Ten different Ittars were procured from the local market based on its popularity and usage. The Ittars were labeled and classified as Methanol and Hexane soluble based on their solubility (Table 1). The Ittars were dissolved in 1 in 10 dilutions for the characteristic study and directly used for animal behavior and biochemical study. All the Ittars and its dilutions were stored and performed in glass vials and glass pipettes respectively.

| Study ID No | Ittar | Solubility |
| --- | --- | --- |
| 1 | Zafran – Zafran (Keesar) | Hexane |
| 2 | Amber – (Pinus Succinifera) | Hexane |
| 3 | Oudh – (Aquilaria agallocha) | Hexane |
| 4 | Muskalkaaba – Deer | Methanol |
| 5 | Woody Rose – Bark of Agarwood | Hexane |
| 6 | Jasmine – Jasmine flower | Hexane |
| 7 | Secret – French perfumes | Hexane |
| 8 | Sandal oil – sandal wood | Methanol |
| 9 | Jannathul Firdous – Wood | Methanol |
| 10 | Unknown Mix | Methanol |

**Table 1:**
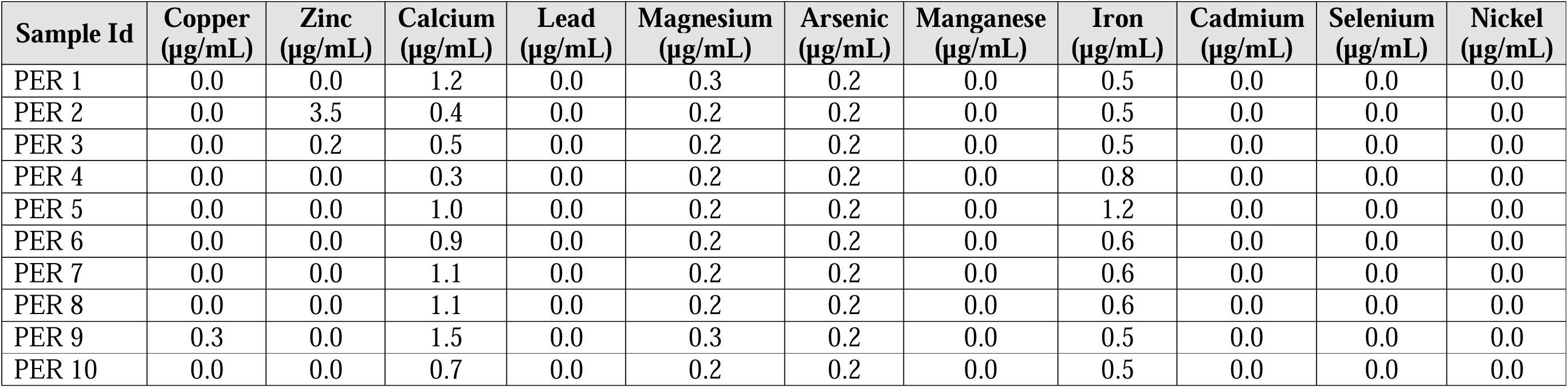
Level of various elements in Ittar based on ICPMS analysis.

### GCMS and ICPMS analysis

The chemical constituents and the elemental composition of the Ittars were analyzed involving GCMS and ICPMS analysis respectively. Diluted Ittar samples were injected and analyzed for its various constituents using the GCMS (Agilent 7890, USA) equipped with split injector and auto-sampler attached to capillary column (Agilent 19091S-43; 30m × 0.25 mm) and mass detector (Agilent 5975C series, USA). Using helium (1ml / min) gas as the mobile phase the sample was separated in a gradient column temperature (35 - 200°C). The obtained mass spectras were acquired in scan mode (70 eV) and analyzed against the standard library. The obtained data were then screened and spectra which has first hit and area more than 0.1% were selected for the study.

The concentration of 9 major heavy and trace elements present in the Ittar samples were analyzed using the quantitative multi-element analysis in NexION 350D (Perkin Elmer, Germany) coupled with ESI-SC2DX auto sampler, cyclonic spray chamber, quartz bore injector and Meinhard concentric nebulizer for sample introduction. The concentration of copper, zinc, calcium, lead, magnesium, arsenic, manganese, iron and nickel were estimated using the respective standard in eight different concentrations which includes PPT, PPB and PPM levels. The level of elements was estimated against the blank and internal standard (Yttrium). GCMS and ICPMS analyses were performed in duplicates.

### Animal experiment

3 months old adult male and female C57BL/6 mice housed separately at 12-12 Light-dark cycle with a body weight ~30 grams were selected for the study. Batches of animals (5 animals per cage) were treated with Ittars for 1, 2 and 3 weeks independently. Male and female mice were treated with Ittar separately in separate battery of cages. The perfumes were applied on the dorsal side of the body below the neck by gentle swabbing. Control animals were remained untreated.

### Behavioral analysis

Control and Ittar perfumed animals were analyzed for its behavior activities such as vertical, horizontal, ambulatory, grooming and rearing using Animal activity meter (Opto-Varimex, USA). The control and perfumed animals were monitored for 2 minutes and recorded for their behavior activities after 1, 2- and 3-weeks post perfume application. The behavior analysis was performed for all the five animals of each set individually and were tabulated for statistical analysis.

### Brain tissue collection

After the behavior study in their respective time points the animals were sacrificed by cervical disc dislocation for collection of cerebral tissue of the nervous system. The cerebral tissue was carefully dissected from the brain using stereoscopic microscope, washed in PBS solution, pooled and stored in liquid nitrogen until use.

### HPLC analysis

The level of Dopamine, 5-HT, 5HIAA and Homovanillic Acid neurotransmitters were analyzed in the pooled cerebral tissue involving High performance liquid chromatographic (HPLC) analysis. 1 gram of pooled cerebral tissue was homogenized using Teflon homogenizer in 0.1 M per chloric acid and 0.05% EDTA buffer for 10 seconds, centrifuged and filtered in 0.45 μm pore filters. Level of neurotransmitter were estimated by performing HPLC in C18 μ-Bondapak Column with a 1.2 ml/ min flow rate and 10-μl-injection volume in Waters HPLC-ECD system (Model No. 2465, Waters Associate Inc. USA). The relative level of the neurotransmitter was analyzed for the perfumed male and female mice against normal mice. All the experiments were performed in triplicates.

### Histological Analysis

The perfume exposed mice skin was carefully removed from the animal using scissors and forceps post hair shaving using auto shaver. The skin samples where fixed in 10% neutral buffered formalin for three days. The skin was then processed using TEK VIP 6 vacuum inflation processor and wax embedded in Bio-optica DP500. Three to five 5micron sections were collected for each whole mount skin sample using microtech cut4050. The sections were stained using Mayer’s Hemotoxylin staining protocol for histological analysis. The slides were further documented using light microscope.

### Data Analysis

All the behavior and expression data obtained from the study were analyzed using Excel and SPSS statistical analysis. The Mean, SEM, Student t-test and statistical significance will be calculated from the data using standard formulas.

### Ethics

All the animal and experimental procedures executed in the work were performed as per the approval of Institutional animal care and ethics committee, Research Center, Prince Sultan Military Medical City, Riyadh, Saudi Arabia (Research protocol No. 30/2015).

## Results and Discussion

### Composition of Ittar

Based on GCMS analysis a total of 392 various chemical constituents were found to be present from all the 10 Ittar samples. Ittar sample 1 and 10 has the lowest constituent of 13 and 6 molecules respectively (Figure 1a). Where as in other Ittar samples 55 (Ittar sample 2), 68 (Ittar Sample 3), 54 (Ittar sample 4), 95 (Ittar sample 5), 72 (Ittar sample 6), 47 (Ittar sample 7), 57 (Ittar sample 8) and 52 (Ittar sample 9) molecules were identified as its major constituents (Figure 1a and 1b). Hexane soluble Ittar showed a total of only two molecules overlapping in all the five Ittars (Figure 1c), whereas only one molecule was found overlapping in all the five-methanol soluble Ittars (Figure 1d). Based on R analysis of the molecule, Ittar 1 and 10 were found to be more similar with Ittar 7, which is related to Ittar 8 and Ittar 9. Ittar 5, 3 and 6 were found distantly related based on molecular components with the other Ittars (Figure 1b).

**Figure 1:**
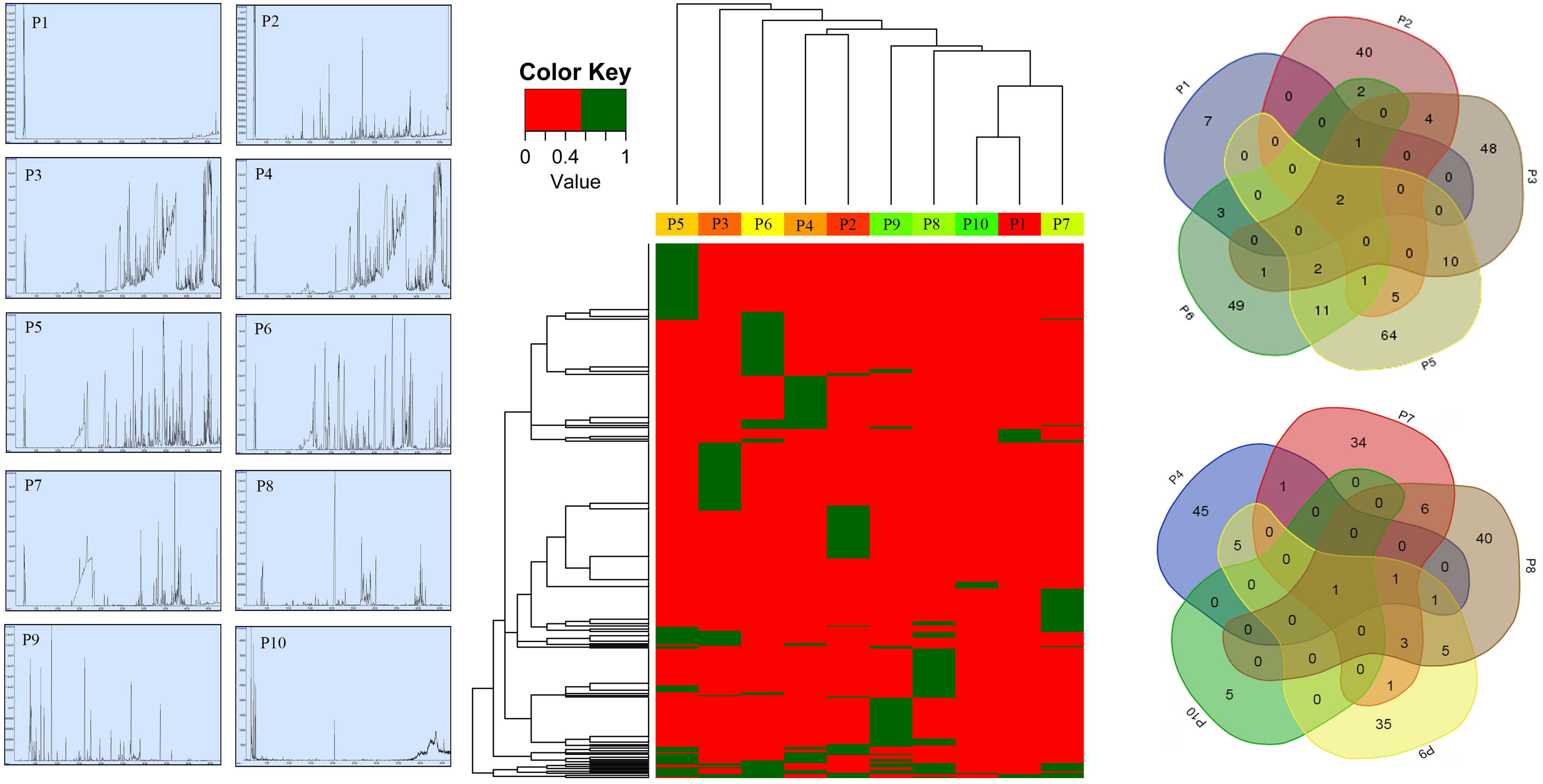
a. GCMS spectra of the 10 Ittar samples; b. Cluster hierarchical analysis of the Ittar samples based on their molecule composition; c. Venn diagram analysis of Hexane soluble Ittar molecules for their molecular content; d. Venn diagram analysis of methanol soluble Ittar molecules for their molecular content.

Based on ICPMS analysis for major heavy and trace elements in the Ittar samples (Table 1) it was found that majority of the elements were either completely absent or present in trace level. Copper, lead, manganese, cadmium, selenium and nickel were found to be completely absent in all the Ittar samples. Zinc metal was not found in all the Ittar samples except in Ittar No. 2 and 4 samples. Calcium, magnesium and iron were found to be present by less than 1.5 PPM (Table 1). No major toxic elements such as arsenic, beryllium, cadmium, lead and mercury were detected at PPM levels in all the perfumes.

### Effect of Behavior

Based on behavior analysis it was found that there was no much obvious changes observed among the perfumed animals in comparison to the non-perfumed animals. Behavioral analysis for horizontal movement showed significant up and down regulation in male mice exposed to Ittar 3, 6 and 7 (Figure 2a). Among female mice Ittar 8 was found effective in altering the horizontal movement. Except for Ittar No.2 in male mice, there was no any significant changes were observed for the vertical movement among both male and female treatment. No significant changes were observed for Ambulatory, Grooming and Rearing were observed among both male and female mice for the perfume application.

**Figure 2:**
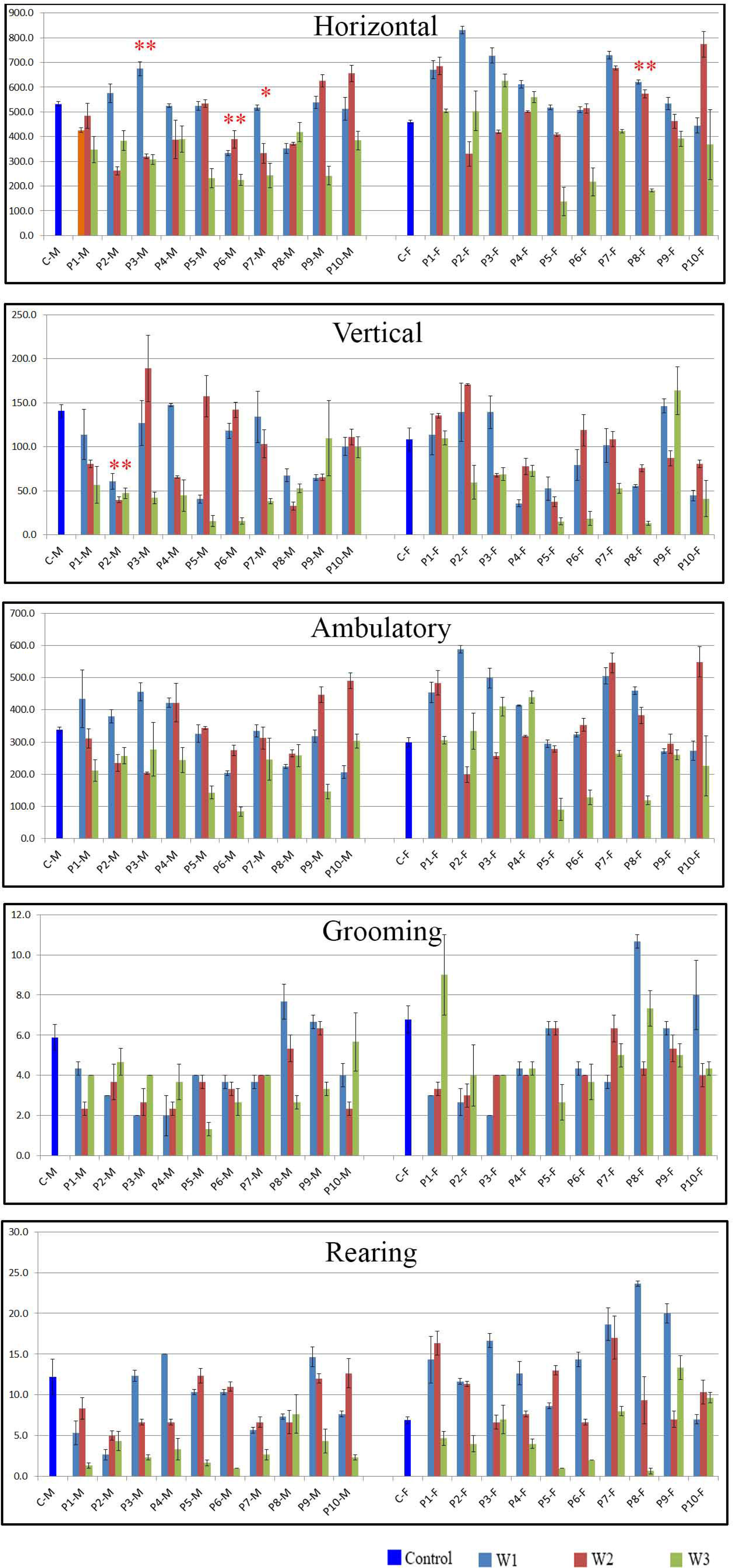
Horizontal, vertical, Ambulator, Grooming and Rearing behavior of the male and female mice treated with 10 Ittar against control animals.

### Neurotransmitter Analysis

A gross up-regulation of neurotransmitter such as Dopamine, 5-HT, 5HIAA and Homovanillic Acid were observed in the brain cerebral tissue upon exposure to Ittar for three weeks. The level of dopamine was non-significantly up-regulated in both the male and female brain exposed to ittar samples 1 to 7 (Figure 3), with no much change in other perfumes. Similarly, serotonin level was also found up-regulated significantly in the same set of perfume exposures. The level of 5-HIAA, HVA and Serotonin were also found elevated non-significantly in all of the exposures (Figure 3).

**Figure 3:**
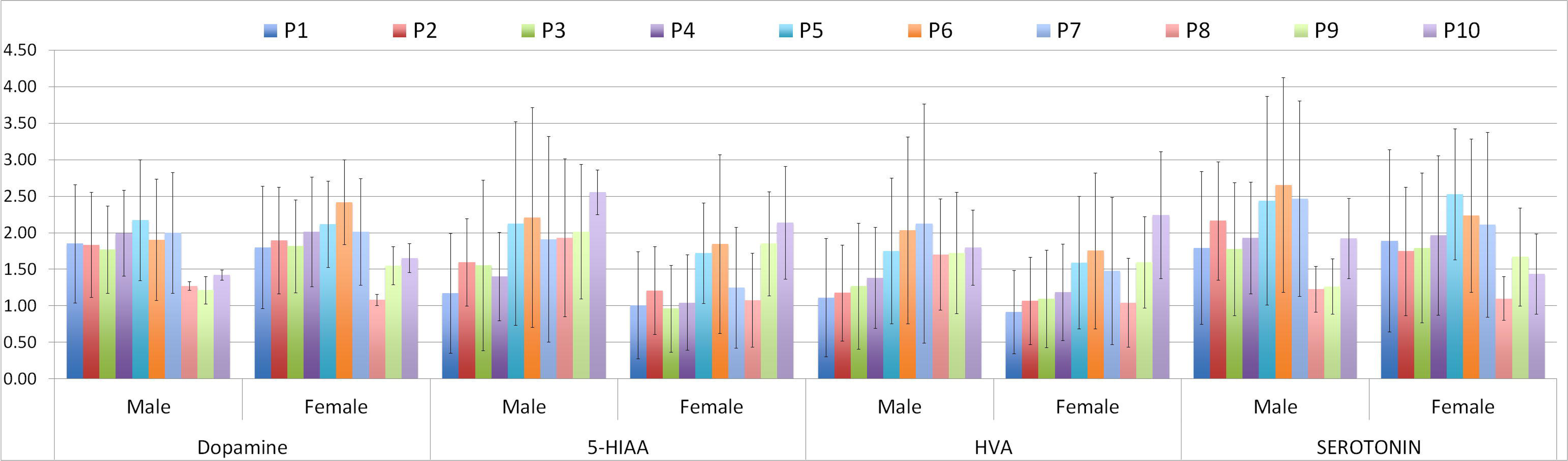
Molecular level of Dopamine, 5-HIAA, HVA and Serotonin in the brain tissue of male and female mice treated with Ittar against control mice.

### Histological Analysis

Histological findings showed a normal pattern of skin epidermal and dermal layers with mild or no infiltration of inflammatory cells in control group of animals. Perfume treated animals showed mild to massive changes in both epidermal and dermal layers characterized by intense infiltration of inflammatory cells, hyperplasia, presence of enlarged and vacuolated cells, hyperchromasia and edema formation (Figure 4 and Table 2). Extreme infiltration of inflammatory cells was observed in perfume 4 and 5 treated skin, followed by 7 and 9 Ittar. Large vacuolated cells were found to be present in skin treated with Ittar 5 and 8 samples (Figure 4 and Table 2). The physiological structure of the skin was compared against the untreated control samples.

**Table 2:** Effect of Ittar in skin samples post treatment for number of epidermal layers, leukocyte infiltration and Edema. (+ = Low; ++ = Moderate; +++ = Extreme).

| Group | Observation |  |  |
| --- | --- | --- | --- |
|  | No. of Epidermal layers | Leukocyte Infiltration | Edema |
| Control Male | 2-3 | - | - |
| Control Female | 2-3 | - | - |
| Perfume 1 - Male | 5-10 | +++ | +++ |
| Perfume 1 - Female | 5-10 | +++ | +++ |
| Perfume 2 - Male | 2-3 | - | - |
| Perfume 2 - Female | 2-3 | - | - |
| Perfume 3 - Male | 2-3 | - | - |
| Perfume 3 - Female | 2-3 | - | - |
| Perfume 4 - Male | 3-8 | ++ | ++ |
| Perfume 4 - Female | 3-8 | ++ | ++ |
| Perfume 5 - Male | 2-4 | ++ | ++ |
| Perfume 5 - Female | 2-4 | ++ | ++ |
| Perfume 6 - Male | 2-3 | - | - |
| Perfume 6 - Female | 2-3 | - | - |
| Perfume 7 - Male | 2-3 | - | - |
| Perfume 7 - Female | 2-3 | - | - |
| Perfume 8 - Male | 2-3 | - | - |
| Perfume 8 - Female | 2-3 | - | - |
| Perfume 9 - Male | 2-3 | - | - |
| Perfume 9 - Female | 2-3 | - | - |
| Perfume 10 - Male | 5-9 | +++ | +++ |
| Perfume 10 - Female | 5-10 | ++ | +++ |

**Figure 4:**
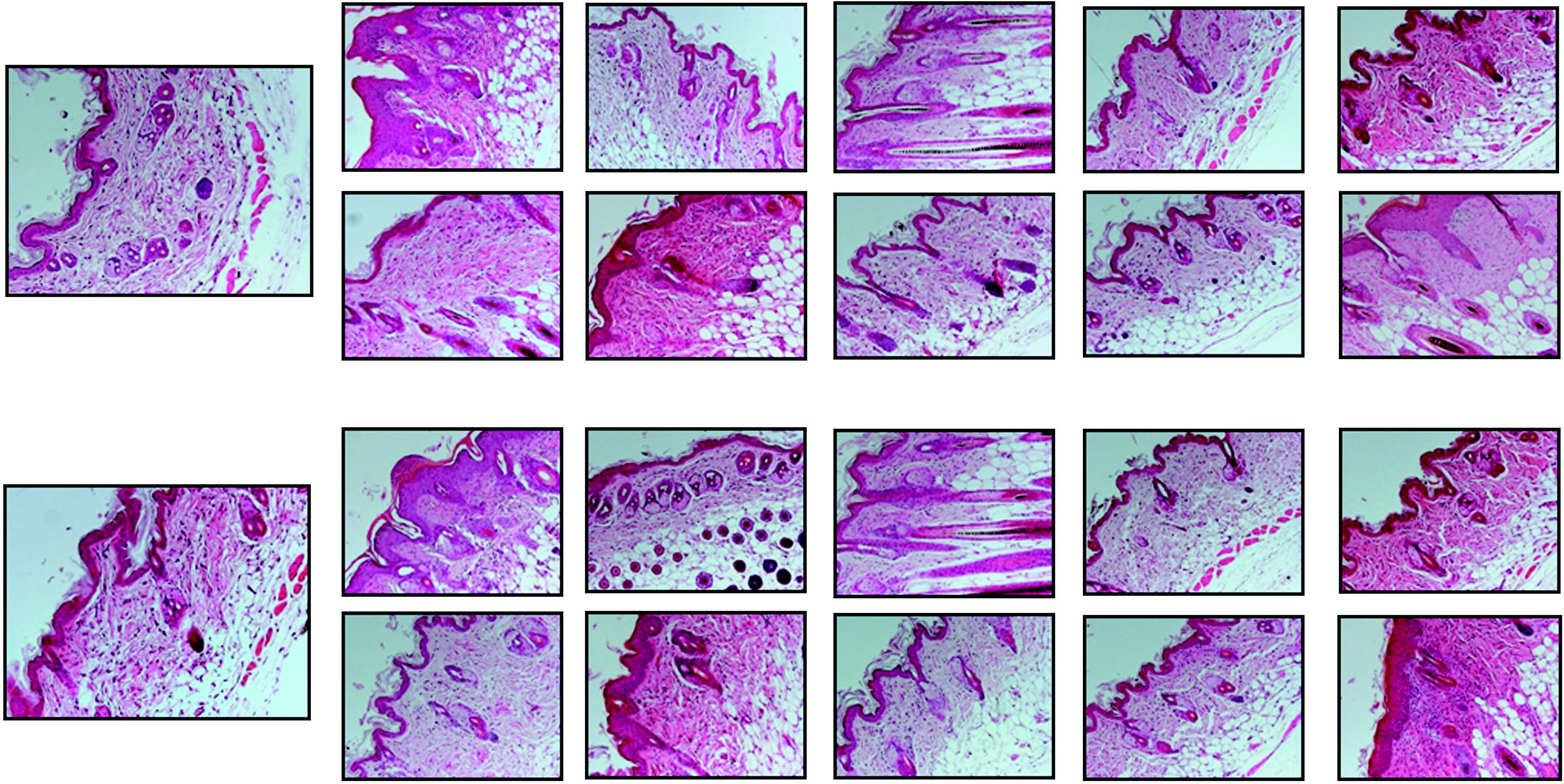
Male and Female mice skin section stained with Hemotoxylin. Panel A details the male skin sections and Panel B details the female skin sections treated with Ittar against control skin section.

## Discussions

Ittars are the traditional non-alcoholic perfume extracted from various flora and fauna through slow distillation. It has been used by mankind for more than thousands of years as a traditional fragrance. Though it is generally called as Ittar, there are more than 100 to 200 different types of Ittars available in the market for its various source and preparation. In this first ever study of understanding the components and effect of Ittar, we have characterized 10 various and widely used Ittar for its composition and its effect on the widely used mammalian model, mice.

Based on this study it is very interesting to see that the 10 different Ittar are completely different for their chemical composition as it is prepared from different plant sources. Apart from the natural small molecules of the plant source, there are several chemical adjuvants been found in the Ittar which have been added during its preparation. The most ubiquitously used Diethyl phthalate (DEP) was found to be present in almost all the Ittar in high index. Known for its wide usage as vehicle for perfumes (Api 2001), DEP has been associated with several toxic reactions such as eye irritation in rabbit (Draize et al 1944); dermal irritation in Albino rabbits (RIFM 1974), penetrates mice, rat and human skin (Scott et al 1987). Presence of DEP in various doses in all the Ittar might have potential toxic effect upon direct exposure to human skin, leading to various dermal related toxicity and infiltration of dermal cells. N-Hexane, Pentane & its various derivative and butane are the major constituents of all the Ittars which has various toxic effects at different doses. Elemental analysis of Ittar ruled out the presence of toxic heavy metals such as arsenic, beryllium, cadmium, lead and mercury.

Behavior of the animals were found to be unaltered with the application of ittar in mice significantly except for the few changes in male and female mice for the horizontal and vertical movement. Thus, this study impacting the fact of no changes in mice normal behavior with the application of Ittar, but the changes for few ittar among male and female mice has got some significance, which has to be studied in-depth. Coarse up-regulation of the neurotransmitter in the brain tissue of mice as effect of perfume is impacting an important association. As the study have been performed on the whole cerebrum tissue, the up-regulation of the transmitter is not significant, a focused antibody staining or biochemical screening on the specific part of the brain tissue might picture an important association of these perfumes in initiating the level of neurotransmitter. The association of perfumes in up-regulating the neurotransmitter can be further studied in stress and Parkinson’s disease model, where the neurotransmitter level goes down, as an alternative reverting factor.

In spite of its several benefits of using Ittar, our study has cautioned a big concern on using Ittar directly on the skin. In this study we have found massive changes in the epidermal and dermal layers of the skin with moderate to severe inflammation, infiltration, hyperplasia and edema upon direct application of the Ittar on the skin. The effect might be due to various chemicals and small molecules present in the perfume. Thus, our study restraints the direct usage of the perfume to the skin, instead it can be applied on the cloth or indirectly.

## Acknowledgements

This work was supported by PSMMC, Riyadh, KSA. The authors are grateful to Noorul Fowzia for critically reviewing the manuscript.

